# Precise measurement of rodent drinking using CLiQR (Capacitive Lick Quantification in Rodents)

**DOI:** 10.64898/2026.03.24.713970

**Authors:** Christopher Parker, Alexis Lam, Ashley Walters, Harrison Carvour, Jack Douglass, Brannen Dyer, Anthony Glorius, Boston Main, Christine Moore, Margaret Niemeier, Avni Patel, Kaladin White, Nicholas Timme

**Affiliations:** University of Cincinnati 2600 Clifton Ave. Cincinnati, OH 45221 USA

**Keywords:** Lickometry, Behavioral Neuroscience, Alcohol Use Disorder, Obesity, Metabolism

## Abstract

Accurate quantification of rodent licking behavior is essential for studies of fluid intake, including investigations of alcohol use disorder and obesity. Lickometry systems vary widely in sensing modality, cost, scalability, and data resolution. Also, many available systems require specialized housing or store only binary lick/no lick data based on thresholding. Here, we present CLiQR (Capacitive Lick Quantification in Rodents), an open-source capacitive lickometry system designed for high-throughput recording of licking behavior in home-cage environments while preserving the full capacitance time series. The system uses MPR121 capacitive sensors connected to custom metal-tipped serological pipette sippers and a centralized desktop computer to record data from up to 24 mice concurrently, with capacity for two-bottle choice experiments. Validation experiments demonstrated that the capacitive signals reliably distinguish licking from non-licking interactions. Total lick counts showed a positive correlation with measured fluid consumption (R^2^ = 0.849, p < 0.0001), confirming that detected events provide a meaningful proxy for intake. All information necessary to reproduce the system is shared openly in this manuscript and online. By combining scalability, full-trace data acquisition, and low cost, CLiQR provides a flexible and extensible platform for high-throughput behavioral neuroscience experiments and enables retrospective improvement of lick-detection algorithms.

## 1. Hardware in context

### Key advantages of the CLiQR system include

- Streamlined procedure with less animal handling, including GUI for ease-of-use by laboratory personnel
- Capacity for recording from 24 mice concurrently, with all data stored for post-hoc analyses
- Low cost when compared with commercial systems

Measuring fluid consumption in rodents is crucial in numerous pre-clinical experimental contexts. For example, alcohol consumption is measured in the study of alcohol use disorder[3–8], while sucrose consumption is measured in the study of obesity[9], motivation, and addiction[10]. The total fluid consumpted by mice is often too small in short-access paradigms to measure easily using changes in fluid weight, necessitating the use of other techniques to measure fluid consumption. In addition, it is often advantageous to record high time resolution licking behavior because licking microstructure carries important information[9,11]. A foundational study by Davis & Smith in 1992 demonstrated that licking behavior is highly structured and informative beyond the intake measure of total volume consumed[11]. More recent reviews of binge and excessive alcohol drinking similarly highlight that patterns of consumption, not only total intake, are key determinants of physiological impact[3,12]. Jeanblanc et al.[12] argue that the defining feature of binge alcohol drinking is not total quantity consumed but the speed and episodic pattern of intake. They emphasize that rapid consumption on presentation of alcohol (referred to as front-loading hereafter) produces higher peak blood ethanol concentrations than the same volume of total consumption over the course of a full experimental trial. Further, Ardinger et al.[3] present a review of front-loading behavior in which they argue that this same pattern of consumption is apparent in both rodent and human studies of binge drinking. The authors hypothesize that this behavior represents a motivational drive for intoxication, which cannot be assumed in cases of slow consumption that doesn’t lead to impairment. This reinforces the necessity of lickometry for measuring the pattern of intake alongside standard measures of intake volume. Therefore, lickometry systems represent key experimental tools in behavioral neuroscience that provide accurate quantification of fluid consumption and licking behavior.

Starting with the “electronic drinkometer” by Hill & Stellar in 1951[13], researchers[14–24] and companies (e.g., Stoelting, Columbus Instruments, LabeoTech, ConductScience) have developed many systems to measure rodent licking behavior. These systems have distinct trade-offs in terms of in cost, ease-of-use, reliability, and data resolution. Lickometry systems fall mainly into three categories: electronic, optical, or capacitive. To motivate our choice of capacitive sensing, we will describe each briefly.

Electronic lickometry systems utilize an electronic circuit with an air gap between a voltage applied to the sipper and conductive flooring in the cage. On licking the sipper, the animal’s body closes the circuit and current passes from the sipper, through the animal, into the flooring, and back to the recording device.

Alongside considerations about physiological effects of electric current, these systems require animal housing that includes a conductive flooring. Unfortunately, typical mouse cages (e.g. Allentown, as our lab uses) are unsuited for this type of system, as they are made of plastic.

Optical lickometry systems utilize light sensors that detect interruptions in a beam of light. These systems use infrared (IR) light and sensor placed directly in front of the sipper. When the animal licks the sipper, the tongue blocks the light beam, and the system registers a lick. To ensure that licks are registered only when there is contact between tongue and sipper, it is necessary to locate the tip of the sipper inside of a recessed enclosure (roughly the size of the animal’s snout). Unfortunately, these systems also necessitate the use of specialized animal enclosures, as standard mouse cages do not have recessed sippers.

Capacitive lickometry systems utilize capacitive touch-sensing circuits to record capacitance values from metal sippers. In these systems, tongue contact with the sipper increases its total capacitance, allowing the system to detect individual licks by using changes in capacitance. These systems necessitate careful considerations about electrical contacts between capacitive sensors and sippers, wire length/placement, and ability of animals to contact the sipper with body parts other than the tongue because they will affect capacitance measurements. However, despite the ability of animals to cause capacitance changes on the sipper without drinking (e.g. by climbing), the different resistances of various tissues combined with the duration of time in contact with the sipper makes it possible to distinguish licking and non-licking behavior by examining the temporal structure of capacitance changes, which requires recording the full capacitance trace. In addition, capacitance-based systems do not require any specific type of housing; they only require that the sipper tip is made of conductive metal and well-connected to the capacitive sensor.

Another consideration we present here is the choice between commercial and open-source systems. In general, commercial systems cost substantially more than open-source systems. For instance, the Stoelting Lickometer System costs $4,995 for a single cage (https://stoeltingco.com/Neuroscience/Stoelting-Lickometer-System~10447) and the LabeoTech Water Dispenser with Lickometer costs $1,000 per device (https://labeotech.com/product/water-dispenser/). In some cases, the increased cost may be justified for the lack of complicated assembly and the promise of customer support.

For our specific needs, we identified and filled a gap: cost-effective lickometry for recording dozens of mice in their home cages concurrently that can save the full capacitive trace, total volume consumed, and perform post-hoc analyses to identify individual licks. Our custom system allows us to continue using the cages provided by the University of Cincinnati Laboratory Animal Medical Services (Allentown NexGen Mouse 500), which allows for reliable assessment of home cage consumption. This is less stressful for the animals, as we do not need to handle them to begin an experimental session (apart from weighing, if desired). Further, developing the system in-house makes future improvements more streamlined.

To further elucidate similarities/differences between CLiQR and other lickometry systems, we present an overview of many available lickometry systems in **Table 1**.

**Table 1.**
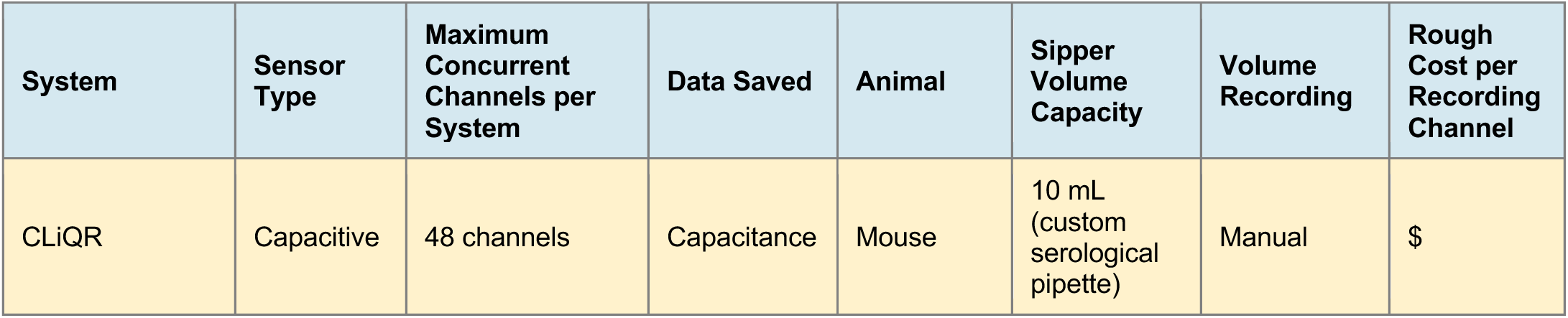
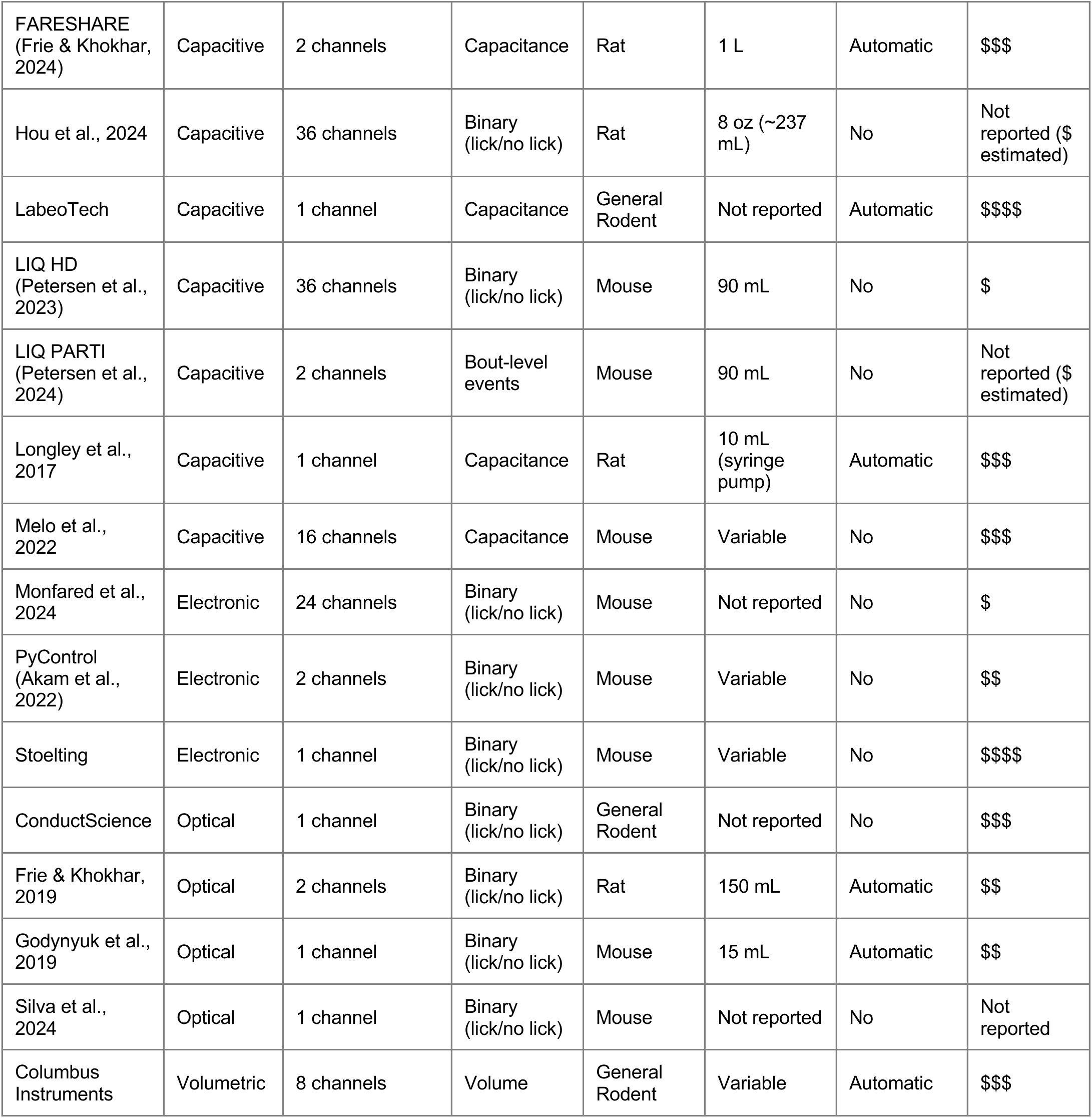
Comparison between various open-source and commercial lickometry systems. The table includes an approximated characterization of the system cost, with the following levels: $ = <$75 per channel, $$ = $75-$150 per channel, $$$ = $150-$300 per channel, $$$$ = >$300 per channel

## 2. Hardware description

The CLiQR system hardware consists of a wire shelving rack with a mounted small-form-factor desktop computer, MPR121 capacitive touch sensors, and 24 custom-built sippers made from serological pipettes with added metal tips (**Figure 1A**).

**Figure 1.**
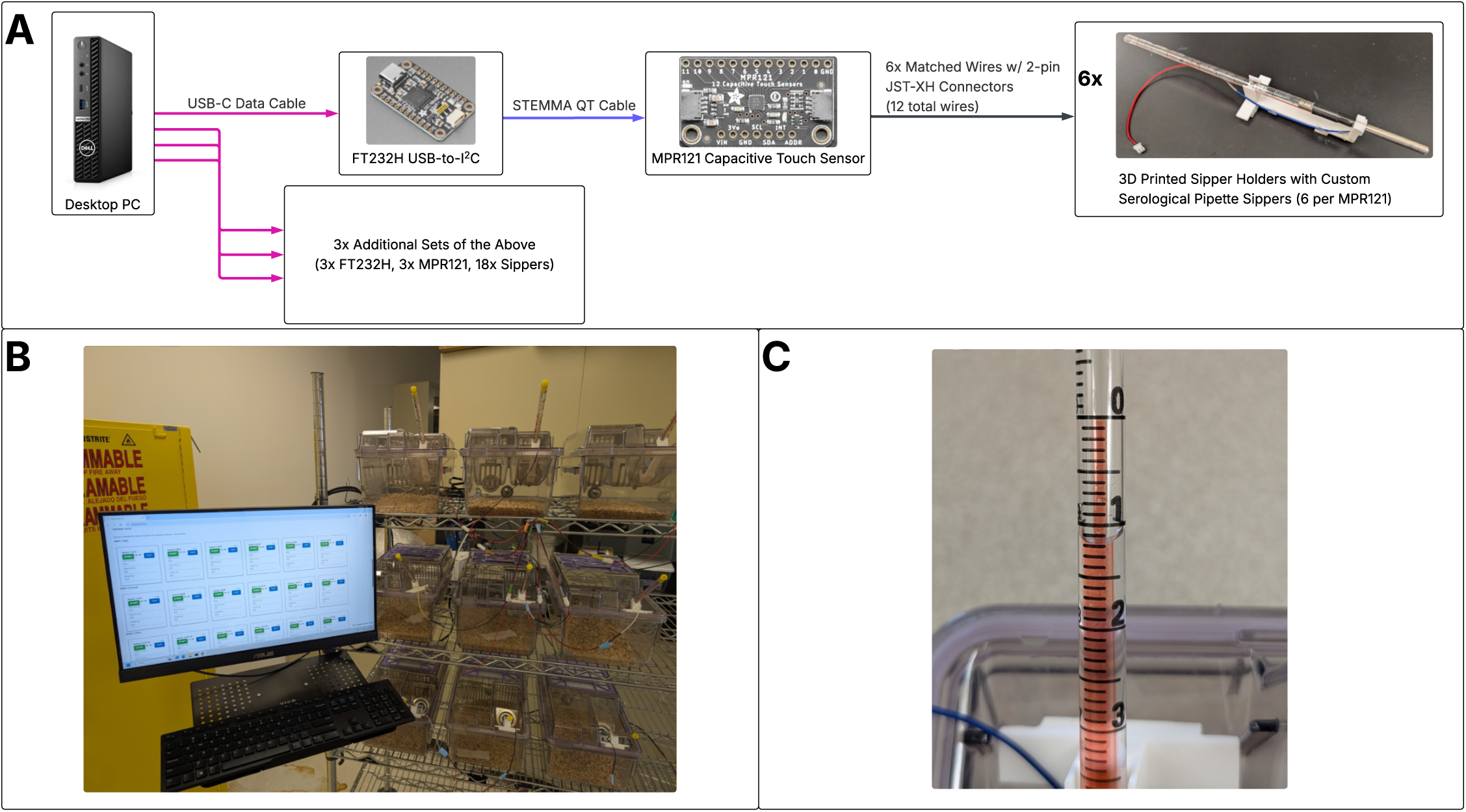
CLiQR System Overview. A) Diagram depicting the system components and connections between them. B) Fully assembled system photograph, showing a portion of the rack with sippers mounted on cages and the mounted monitor displaying the CLiQR GUI. C) Close-up of one of the custom sippers made from serological pipettes. This photograph shows a sipper loaded with 8.8 mL of water (0 = 10 mL, 1 = 9 mL, etc.).

Each MPR121 board has 12 capacitive channels, and we have allocated 2 channels per animal. This future-proofs the system for 2-bottle choice experiments, although currently half of the channels are unused.

The desktop computer runs a Python-based GUI (**Figure 1B**), and post-hoc processing of capacitive data is performed with a separate script (available in MATLAB or Python). Our system uses a Dell Optiplex 7080 running Windows 11 Enterprise, but other small-form-factor desktop computers running other operating systems could be used.

The custom sippers, made from serological pipettes, include graduation markings at 0.1 mL intervals (**Figure 1C**). At the end of a session, the system calculates the difference in recorded volumes to obtain a measurement of volume consumed.

The system saves data using the HDF5 format; a single file contains the full capacitive trace for all 24 animals. The file includes metadata indicating the start and end volumes recorded from the graduated sippers and the weight of the animals. The system starts and stops recording from all animals at once but allows the user to record start and stop timestamps for each animal (used to trim the data to the portion of interest). Our recordings last for 2 hours and sample at ∼110 Hz, with the full data file requiring ∼375 MB of memory.

Data analysis is performed with a modest improvement over single-number thresholding procedures. First, given the MPR121s’ discrete 8-bit integer outputs, we extract the set of all unique values in the recorded traces. Threshold values are computed as the midpoints between each pair of adjacent unique values, yielding N−1 candidate thresholds for a trace containing N unique values. The algorithm then evaluates each candidate threshold in turn, from smallest to largest. For each threshold, all contiguous segments of the trace below it are identified as candidate lick events. Two rejection criteria were applied: (1) segments lasting >167 ms were discarded for exceeding the maximum plausible lick duration in mice (average lick duration is 100-125 ms); (2) any lick candidate in which no sample falls below the threshold located twenty positions lower in the ranked threshold list is discarded, ensuring that shallow deflections corresponding to noise are not counted. After evaluating all candidate thresholds, the threshold yielding the largest number of licks is selected. For more information on the algorithm, we direct the reader to data_analysis.py in the provided code repository (https://github.com/TimmeLab/CLiQR/blob/main/data_analysis.py or https://data.mendeley.com/datasets/wpskw7k3mt/1/files/3d83a2ef-4380-48ed-beb6-d3e78b43fa8d).

We declare here that Claude Code (Anthropic; primarily Opus 4.8) was used to generate significant portions of the codebase for this project. Development used the Superpowers[25] plugin to enforce more thorough planning and test-driven development. All code was reviewed by the authors, and we take full responsibility for the final code.

## 3. Design files summary

Up-to-date versions of the code, 3D print model files, and documentation are available on our lab GitHub: https://github.com/TimmeLab/CLiQR. We have also uploaded all supplemental materials (including additional validation data that is not uploaded to GitHub) to Mendeley Data: https://data.mendeley.com/datasets/wpskw7k3mt/1. The design files summary table uses the filenames found on Mendeley Data (as the system is improved over time, the GitHub will potentially change).

| Category | Design file name | File type | Open source license | Location of the file |
| --- | --- | --- | --- | --- |
| Sipper Holders | AllentownSipperHolderRingContact.stl | 3D Print File | CC BY 4.0 |  |
| Rack Mounts | FTBoardMount.stl<br>MPR121_mount_with_cutouts.stl | 3D Print Files | CC BY 4.0 |  |
| Software | Numerous, highlights include:<br>recording_gui.py<br>DataAnalysis.ipynb<br>data_analysis.py<br>components/<br>recording/<br>hardware/<br>utils/ | Software | MIT |  |

### Design file details by category

#### Sipper Holders

- AllentownSipperHolderRingContact.stl is a 3D print file for the plastic holders that connect the sippers to the capacitive sensing boards. See below for instructions on assembly after printing.

#### Rack Mounts

- FTBoardMount.stl is a simple mount for attaching the FT232H boards to a wire rack.
- MPR121_mount_with_cutouts.stl is a similar mount for attaching MPR121 boards to a wire rack.

#### Software

Brief descriptions of the notable software files follow (this list is not exhaustive):

- recording_gui.py is the entry point for the Solara-based Python GUI that runs the system.
- DataAnalysis.ipynb is the Jupyter notebook that calls functions in data_analysis.py for data analysis and visualization. The Jupyter notebook exports data summaries in various CSV files for plotting in Prism or other software.
- data_analysis.py contains the functions used in the DataAnalysis.ipynb notebook.
- components/ is a sub-module that includes all of the various Solara GUI components used in recording_gui.py
- recording/ is a sub-module that includes code for asynchronous recording
- hardware/ is a sub-module that includes code for communication with the FT232H and MPR121 boards
- utils/ includes state.py which stores the persistent state of the Solara GUI between runs. This is where things like the FT232H board serial to MPR121 sensor number mapping live.

## 4. Bill of materials summary

See below for a partial bill of materials for the CLiQR system. We have included the major costs for the system in this table. See the full bill of materials on Mendeley Data (https://data.mendeley.com/datasets/wpskw7k3mt/1/files/56b3c5b3-4b72-4035-b960-8ed6595181f0), which includes all incidentals and consumables also.

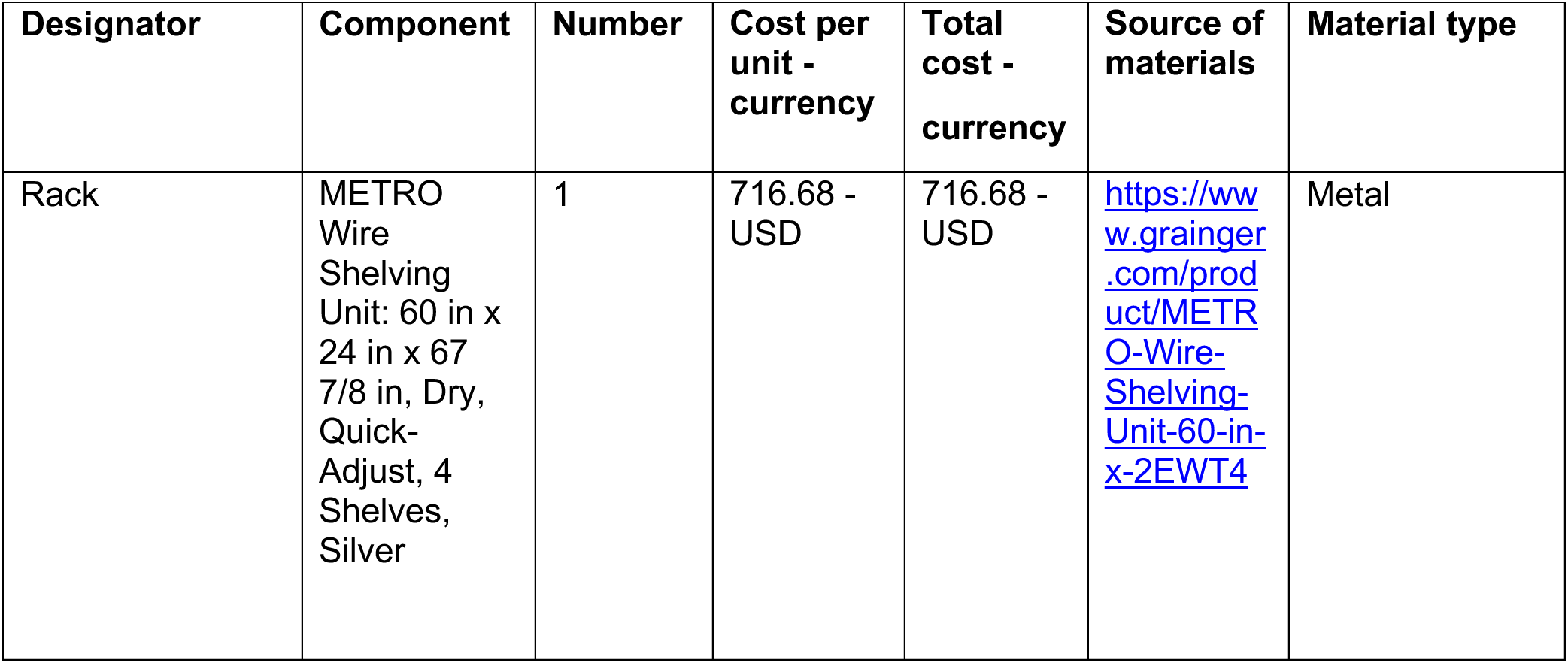
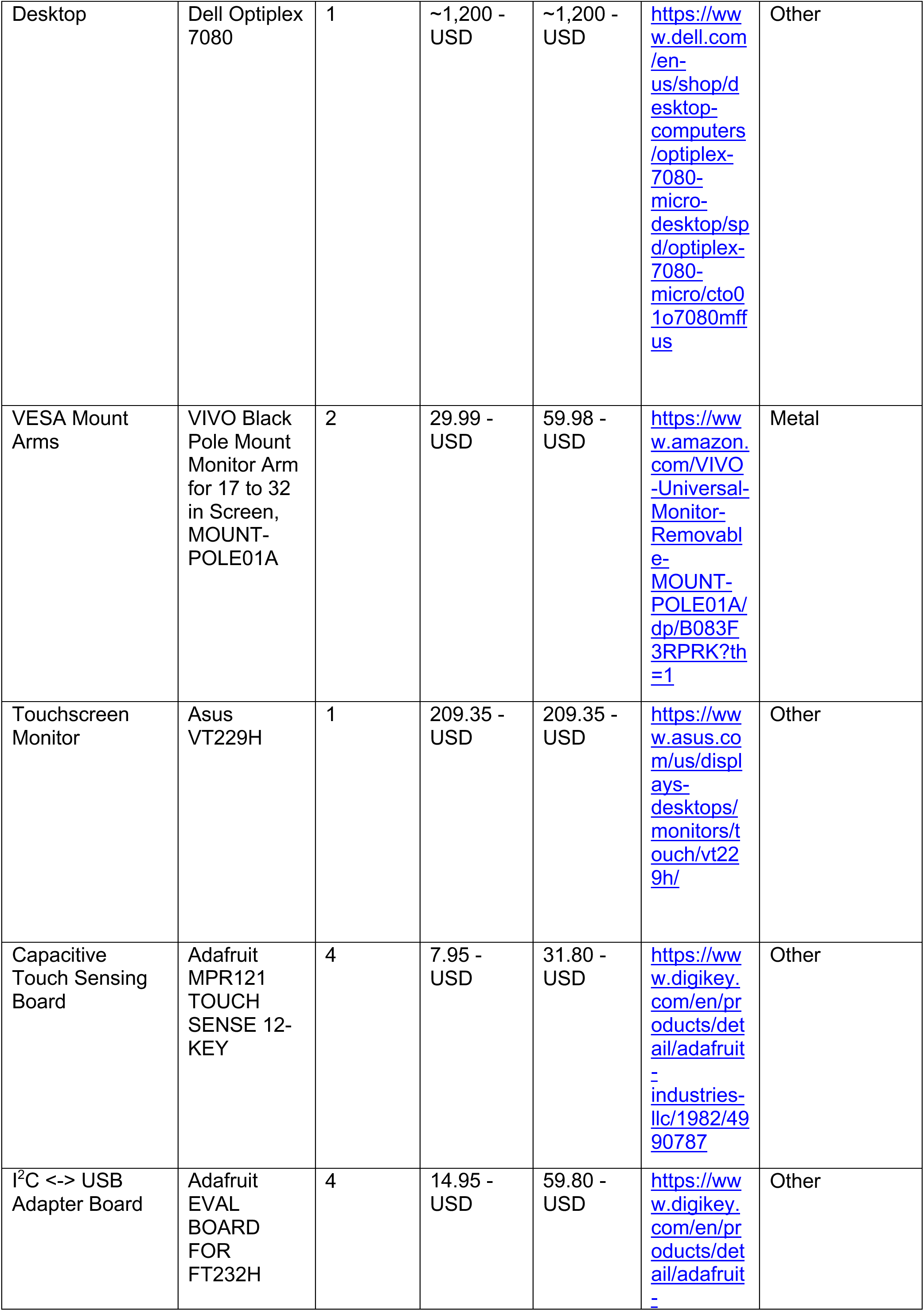
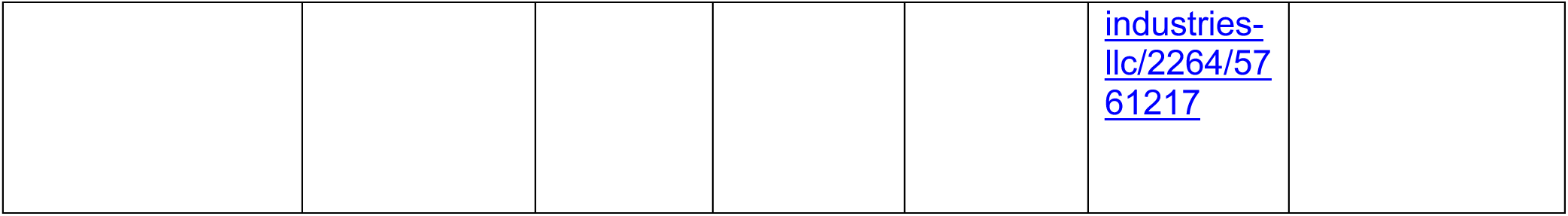

The desktop computer that we use was given to us by our department free of charge. The exact desktop that we use is no longer available from Dell (Optiplex 7080 with an i7), but the system can be run on a less powerful machine. The Dell Optiplex 7080 with an i5 is currently 989 USD and represents a suitable alternative option.

## 5. Build instructions

This section will include step-by-step assembly instructions and figures depicting these steps. These instructions can be found with additional photographs in the Supplemental Data (https://data.mendeley.com/datasets/wpskw7k3mt/1/files/3e83738e-b9dd-4ce7-99c1-0470d7cf241e).

B1. Assemble the wire rack and attach the desktop computer and a power strip at the back of the top shelf with zip ties (**Figure 2A**). Our shelving rack was designed with Imperial units, and the shelves are 12 inches apart (∼30.5 cm). We find that this spacing leaves a reasonable amount of headroom over Allentown NexGen Mouse 500 cages.

**Figure 2.**
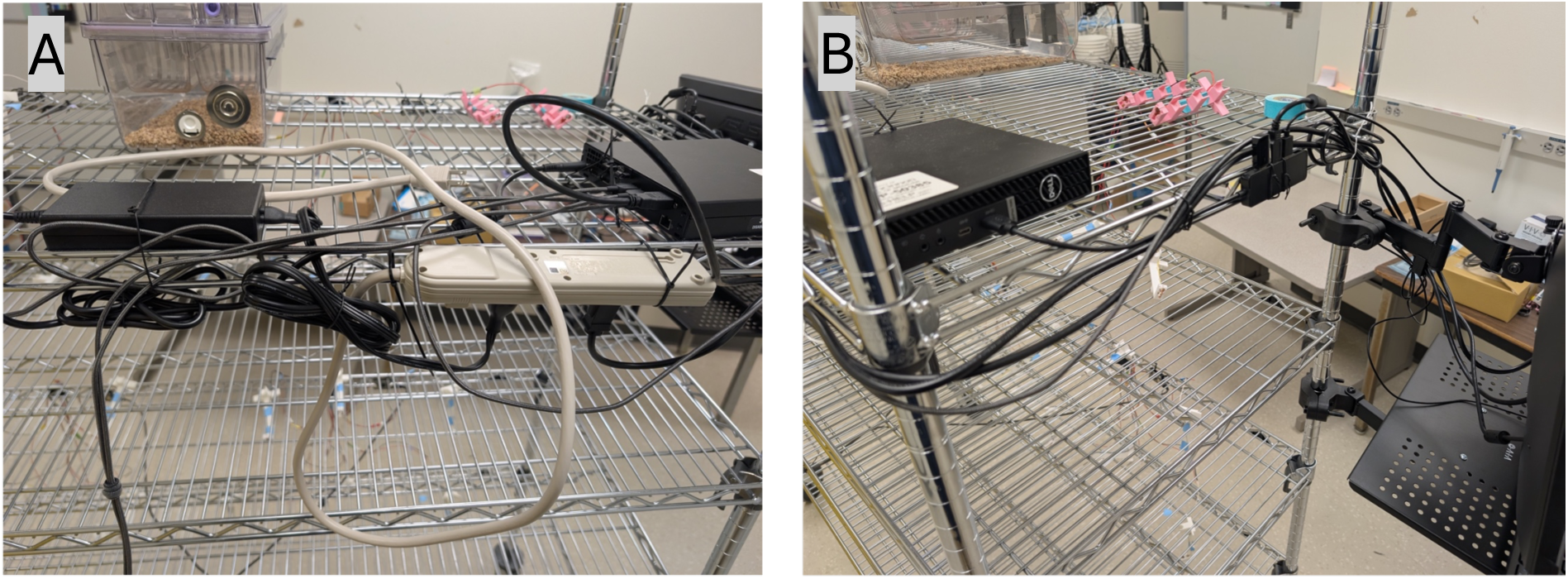
A) View from behind and above the rack showing desktop mounting position. B) Side view of rack, showing desktop, USB hub, and VESA mounting arms.

B2. Attach two VESA mounting arms to the rack and use them to mount a touchscreen monitor and a keyboard tray (see **Figure 2B** for a side view of the rack that shows the desktop, USB hub, and VESA mounting arm positions).

B3. All peripherals (keyboard, touchscreen touch function, WiFi adapter) should be attached to the PC with a USB hub (**Figure 2B**).

B4. Assemble and attach the capacitive sensing chips to the system. The following photographs show the assembly of the breakout boards with their 3D printed mounts and how each component is wired to the others.

a. To begin, we show a single FT232H board mount in **Figure 3A**, and the same mount with an FT232H attached in **Figure 3B**.
b. Before attaching the MPR121 sensors to the 3D printed mounts, wires need to be soldered to the capacitive sensing pads. As shown in **Figure 4A**, we attach the wires so that they extend from the back of the board.
c. Next, we show a single MPR121 board mount in **Figure 4B** and the same mount with an MPR121 attached in **Figure 4C**. Note that the 3D printed mounts for these boards differ
d. somewhat, as the MPR121 is only secured with mounting screws on one side. Both designs utilize M2.5 heat set threaded inserts to ensure secure mounting with M2.5 screws.
e. Note the liberal application of hot glue to the board and mount where the wires attach to the board (**Figure 4C**). This not only serves to further fasten the board to the mount but also helps prevent some mechanical strain on the wire connections from repeated bending when placing sippers on cages.
f. For the wires attached to the capacitive sensing pads, we use JST XH 2-pin connectors. The connectors allow replacement of broken sipper holders without soldering (the pads are hidden under a thick layer of hot glue, making resoldering a hassle). Currently, only one of the two wires is used and the other is future-proofing in case we decide to do two-bottle choice experiments.

**Figure 3.**
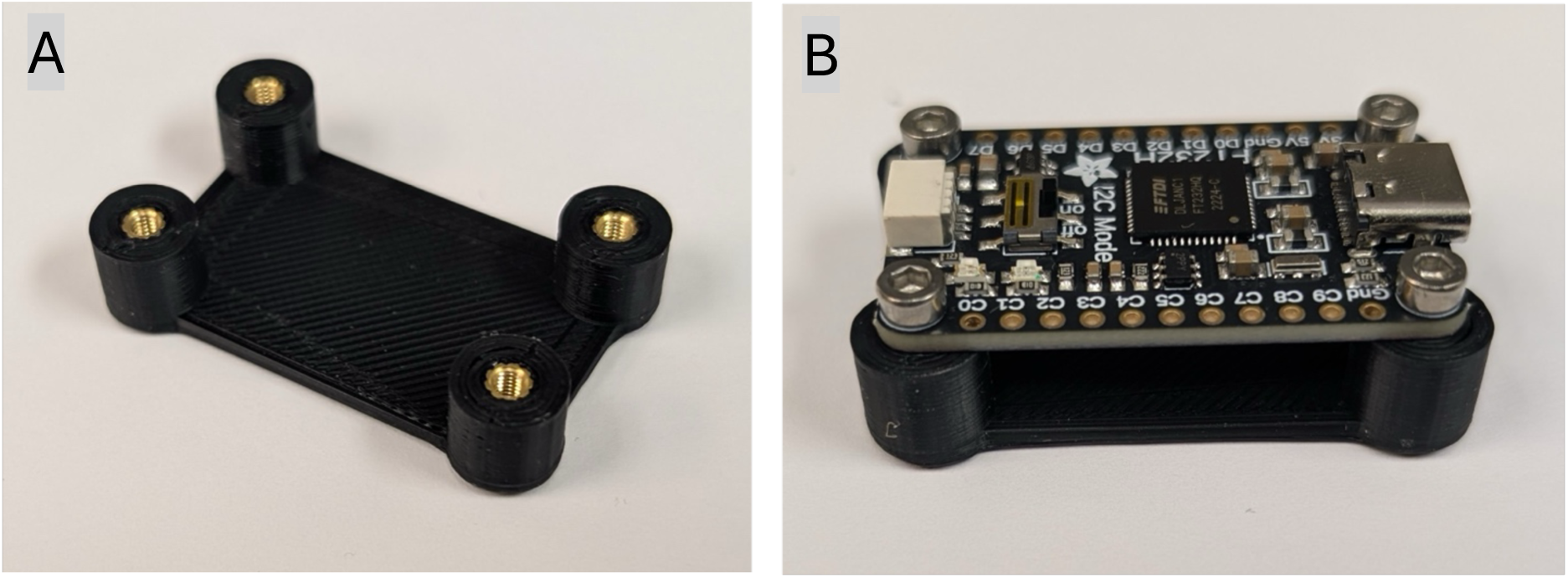
A) 3D Printed mount for FT232H breakout board. The STL file for this print is FTBoardMount.stl. Note the presence of M2.5 heat set threaded inserts for mounting screws. B) FT232H Board attached to 3D printed mount with M2.5 screws. Note the USB-C connector on the right and the STEMMA QT connector on the left.

**Figure 4.**
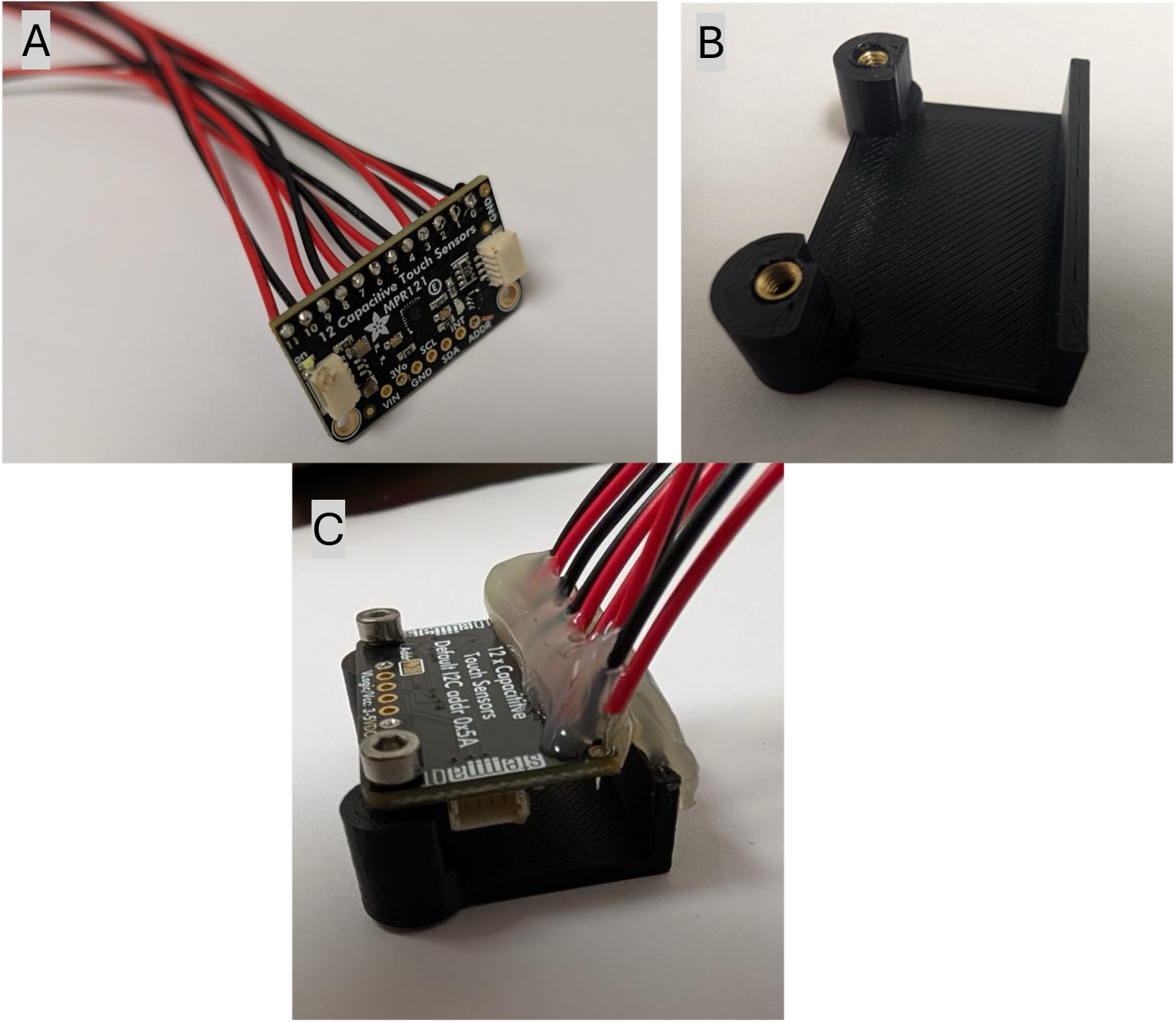
A) MPR121 with wires attached to capacitive sensing pads. Note the orientation of the board with the wires extending from the rear. B) 3D Printed mount for MPR121 capacitive sensing board. Also uses M2.5 heat set threaded inserts for mounting screws, as with the FT232H mount. STL file is MPR121_mount_with_cutouts.stl. C) MPR121 board attached to 3D printed mount. Hot glue is applied liberally to the wires and joins the edge of the board to the mount. This is necessary to ensure mechanical strain doesn’t cause wires to detach.

B5. The next hurdle is making good electrical contact between the MPR121 and a metal sipper tip. For this purpose, we designed custom sippers and 3D printed sipper holders. See **Figure 5A** for a photograph of the instructions for creating the custom sippers and **Figure 5B** for an assembled sipper.

**Figure 5.**
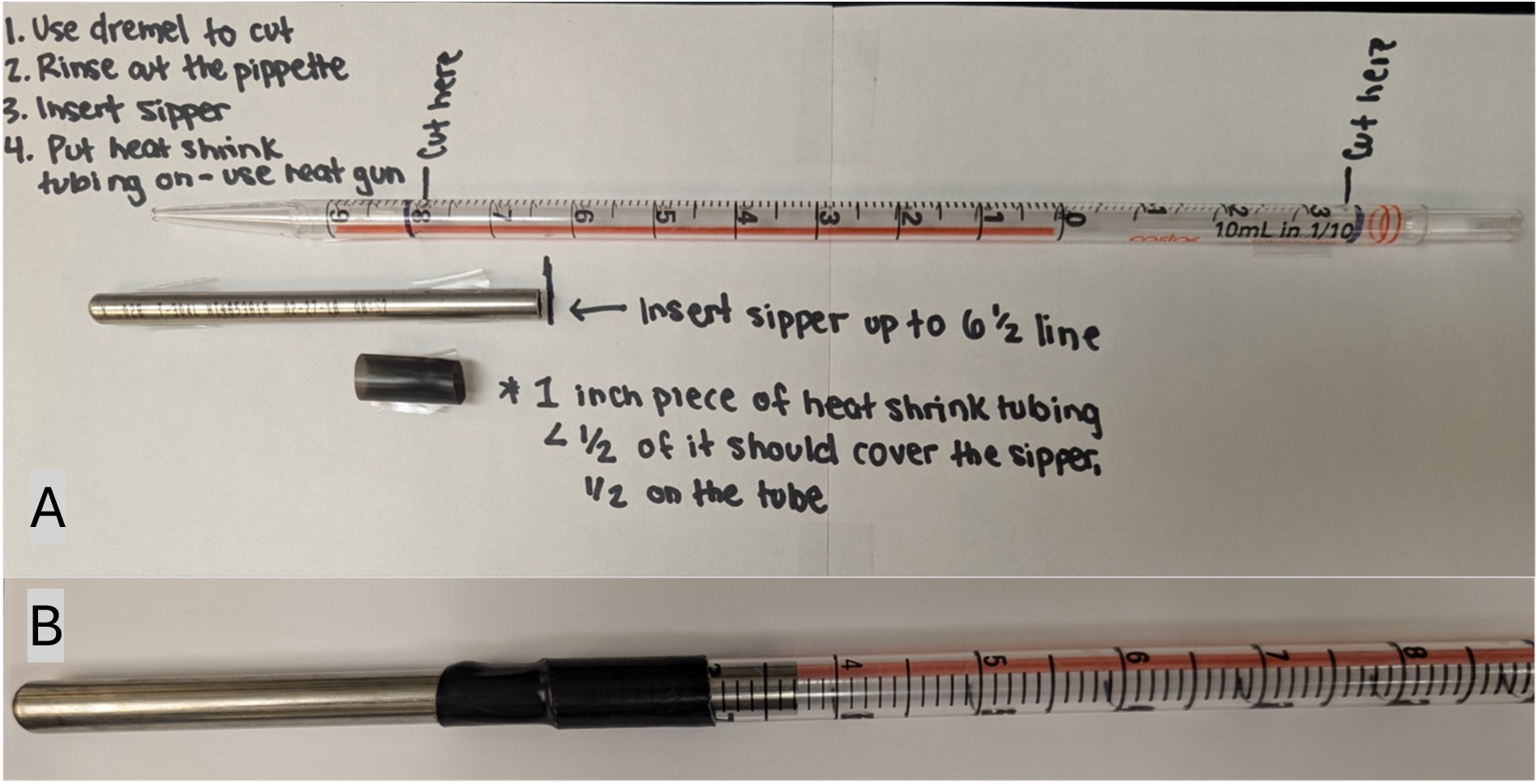
A) Instructions for cutting plastic serological pipettes to size and adding metal sipper tip (attached with heat shrink tubing). B) Custom sipper made from serological pipette and metal tip.

B6. The 3D printed sipper holders are designed with a loop of copper tape through which we push the metal sipper tip to ensure solid electrical contact. **Figure 6A** shows a sipper holder before wiring. White filament is used for printing these to ensure that reading from the graduated sippers is possible (dark filaments make them difficult to read in the dim light of the vivarium night cycle).

**Figure 6.**
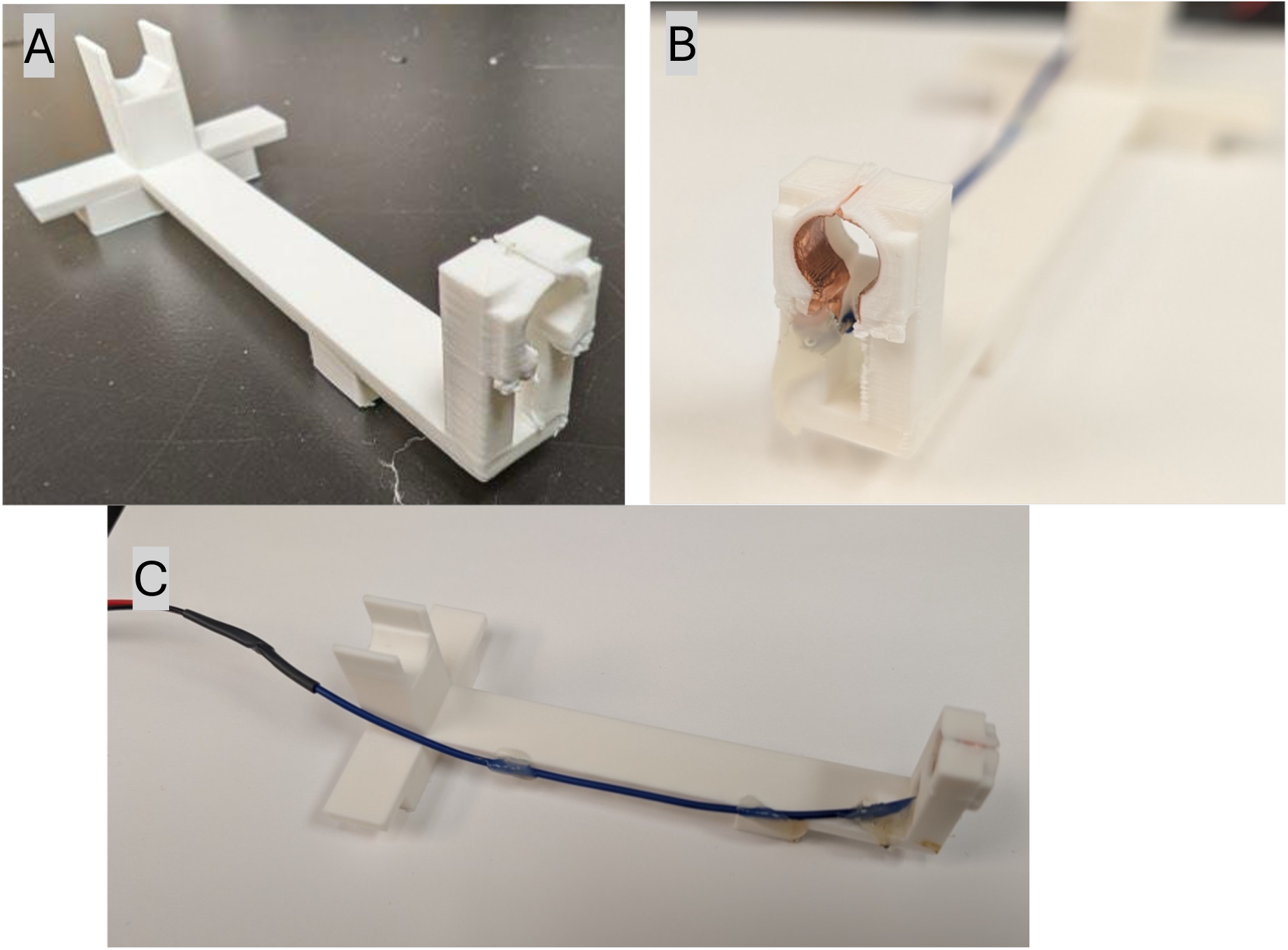
A) 3D printed part before wiring. B) Ring contact with copper tape and wire. The wire is soldered to the copper tape, and the contact is coated with hot glue to avoid tearing the tape if the wire is yanked. C) Side view of the assembled sipper holder. The wire is hot glued at several points along the side of the sipper holder to further ensure that the wire will not detach easily.

B7. **Figure 6B** shows a sipper holder after having copper tape applied to the inside of the contact loop and a wire soldered to the tape. Hot glue is used to thoroughly coat this connection point to ensure that the tape does not rip during handling of the wire.

B8. **Figure 6C** is a side view of a fully assembled sipper holder showing the additional hot glue applied to the wire at several points along the length of the holder, further securing it and making accidental detachment of the wire less likely. To ensure the wire is long enough to reach from the MPR121 to the cages, we use a separate length of wire for the sipper holder. This wire is then soldered to the JST XH 2-pin connector wire and the joint is covered with heat shrink tubing.

B9. The final setup requires the 4 pairs of FT232H/MPR121 boards to be attached to the rack and wired to the desktop. **Figure 7** shows the relative locations of one pair of boards attached to the rack.

**Figure 7.**
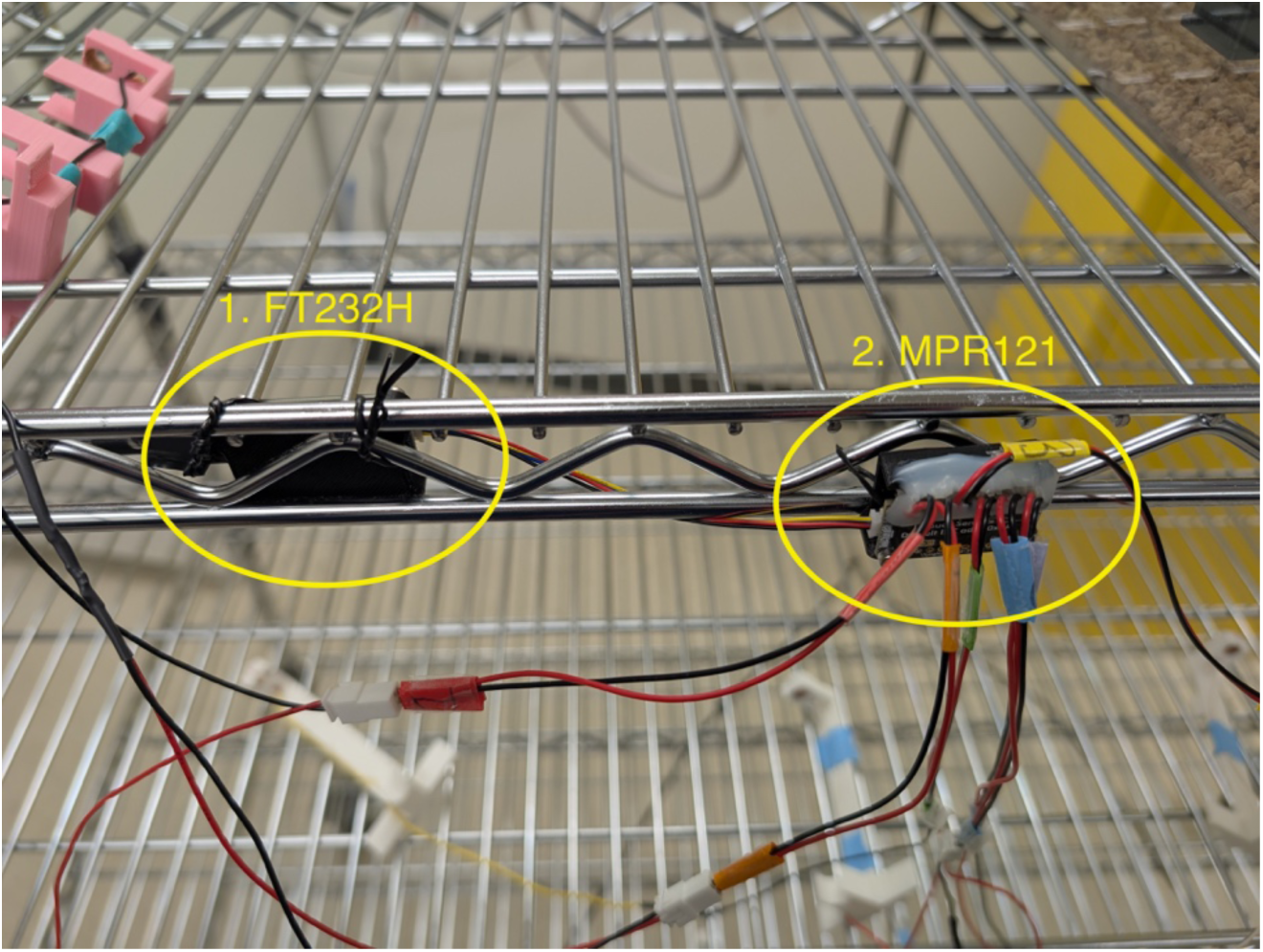
Attachment of 1) FT232H and 2) MPR121 boards to the wire rack. Note the use of colored tape to label each sensor lead to avoid mixing up sensors when placing on cages.

Following the completion of hardware setup, the software will need to be downloaded and configured. Software installation instructions follow:

B10. Windows Driver Setup.

a. Before first use on Windows, you must install drivers for the FT232H boards using Zadig:

i. Download Zadig from https://zadig.akeo.ie/
ii. Follow the steps outlined here: https://learn.adafruit.com/circuitpython-on-any-computer-with-ft232h/windows

B11. Software Download

a. Download the code repository from Mendeley Data (https://data.mendeley.com/datasets/wpskw7k3mt/1/files/7b1ceb77-a009-4f93-8ade-e969073be1cc) and extract the .zip file.
b. Alternative: Clone https://github.com/TimmeLab/CLiQR (or download and extract the .zip file).

B12. Python Installation and Configuration

a. Download and install Miniforge: https://github.com/conda-forge/miniforge/releases
b. On Windows, use the “Miniforge Prompt” from the Start menu. On Mac/Linux, use your normal terminal. Then run:

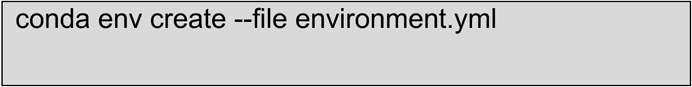

c. On Windows: Hold Shift and right-click start_cliqr.bat, then select Send To -> Desktop (create shortcut). You can then rename the desktop shortcut as you wish, I have just called it “CLiQR.” Double-click the shortcut to launch the GUI without using the command line.
d. Otherwise, you can start the system with the following commands:

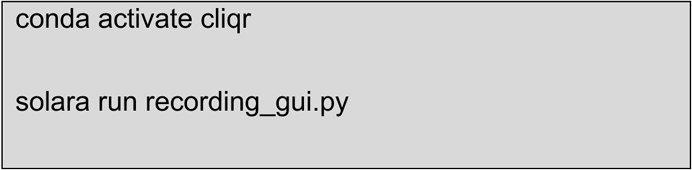

e. The GUI will open in your default web browser at http://localhost:8765

B13. Hardware Configuration

a. The FT232H boards must be assigned serial numbers so the system can identify them consistently, even if the USB ports end up being shuffled somehow. Connect each FT232H board one at a time and run:

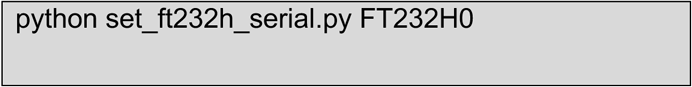

b. Repeat for each board, using serial numbers FT232H0 through FT232H3. These serial numbers map to sensors 1-24 as defined in utils/state.py. If you use different serial numbers, update the SERIAL_NUMBER_SENSOR_MAP constant.

## 6. Operation instructions

These instructions have been adapted from our lab protocol document for running the trials. The full document with additional screenshots can be found in the Supplemental Materials (https://data.mendeley.com/datasets/wpskw7k3mt/1/files/06531472-ce95-4b1b-8cca-6c28c53ff0d2).

O1. The first step should be filling the graduated sippers with your fluid of choice so that it’s easier to quickly start the animals drinking in sequence (without long gaps between start times).

a. Fill the sippers with ∼10 mL and ensure that the rubber stoppers are firmly inserted.
b. Filling should be done with a finger or palm covering the tip of the sipper, because the liquid will quickly leak out otherwise.
c. Once the rubber stopper is inserted, ensure that the sipper is not leaking excessively by uncovering the tip while holding the sipper over a sink or the drain in the floor.
d. If the sipper leaks (more than 1 or 2 drops), double check the stopper. If that doesn’t solve the issue, set that sipper aside.

O2. Double-click on the desktop shortcut with the name “CLiQR” and the recording interface should display after a few seconds. At this point, your screen should look roughly like **Figure 8**.

**Figure 8.**
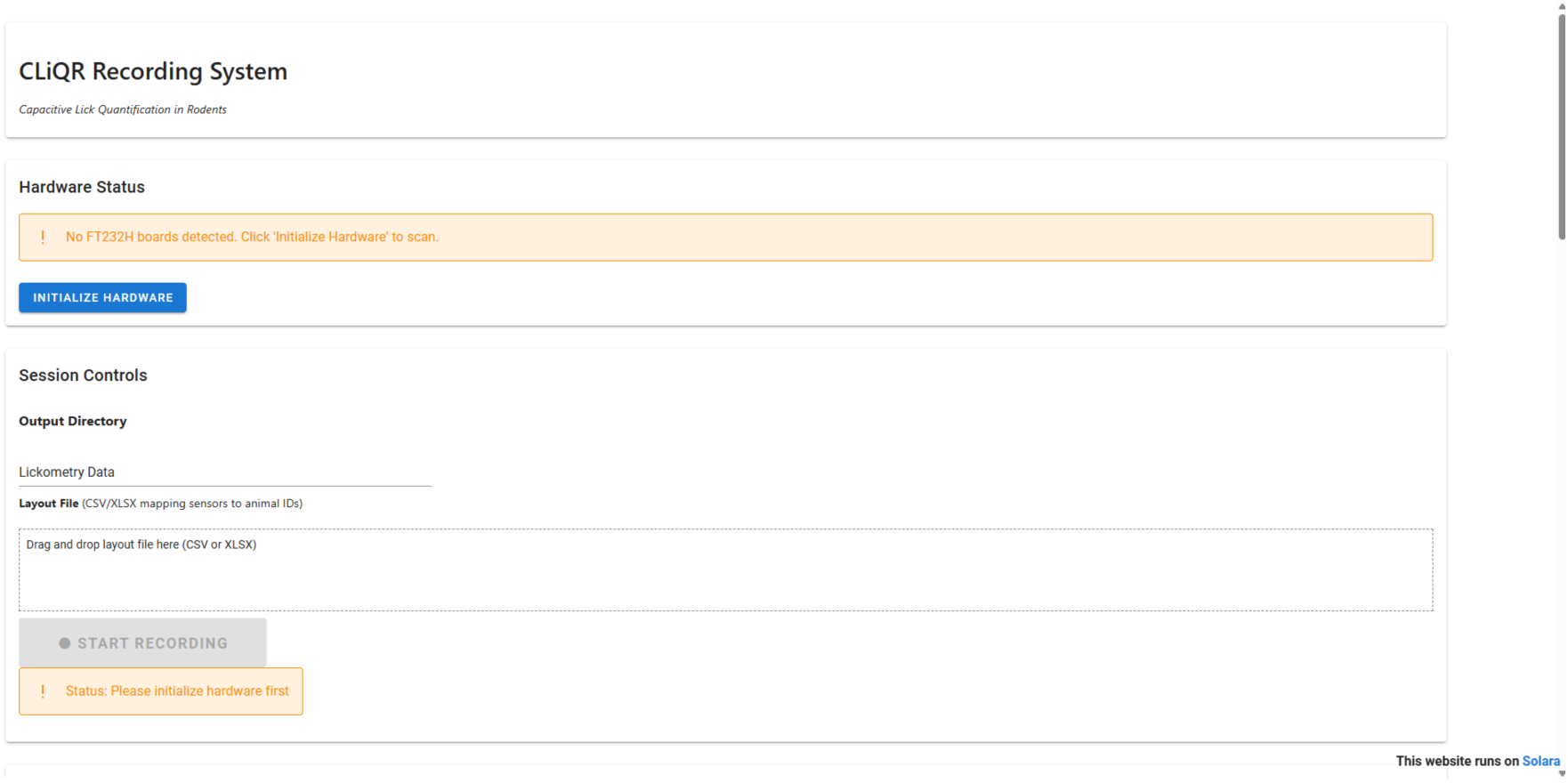
Initial CLiQR GUI window appearance.

O3. Begin by pressing the “Initialize Hardware” button, and the system will detect and configure all of the capacitive sensors.

O4. The “Output Directory” text entry box defaults to “Lickometry Data” (a subdirectory inside the CLiQR directory, just for storing data recordings)

a. This can be changed to include the cohort identifier, like “Lickometry Data/AEW6”, which will create a subdirectory for that cohort inside of the lickometry data subdir.

O5. The “Layout File” section is for assigning animals to sensors so that it stays consistent between drinking sessions. On the first session of a cohort, this will need to be created (and it will be reused each day).

a. This can be created in Excel or just a text editor.
b. The format should be sensor_number, animal_id and each sensor should be on a new line.
c. For instance, 1,AEW6-1 shows the animal ID on sensor 1.
d. Ensure the file is saved as csv and drag-and-drop it onto the cage layout upload box in the CLiQR GUI.
e. The resulting screen should look like **Figure 9**.

**Figure 9.**
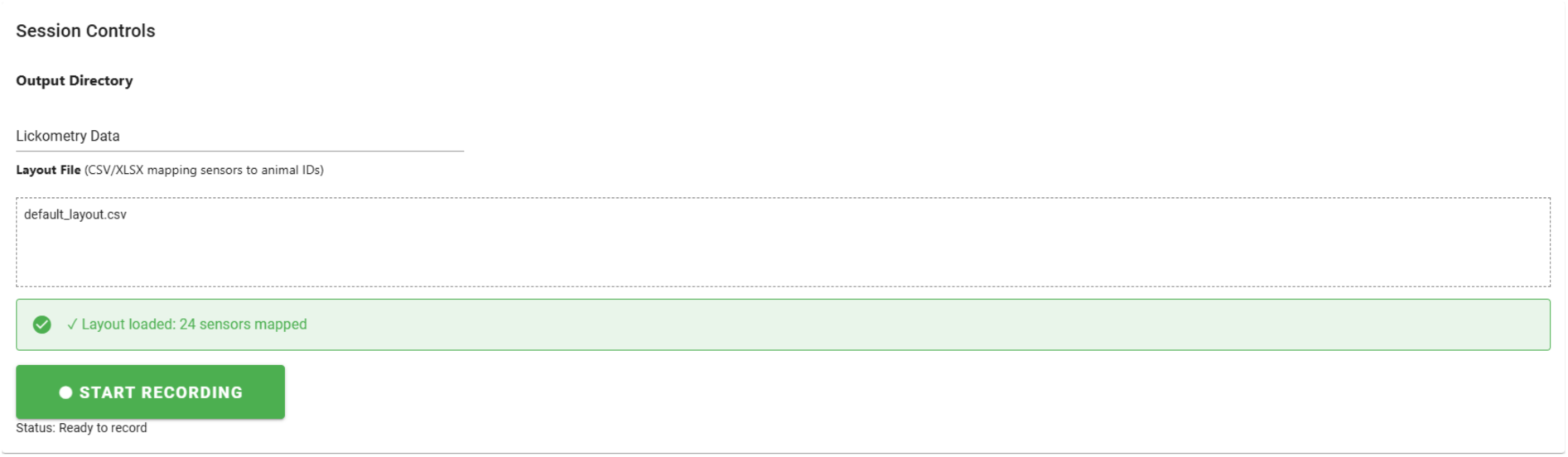
Layout file upload confirmation.

O6. If not already done earlier in the day, weigh each animal using the scale and beaker

O7. Record the animal weights (whether directly measured just now, or from the data sheet with animal weights from earlier) in the “Weight” input box corresponding to the correct sensor for each animal.

a. Now, you should place the animal cages on the rack (6 cages per shelf). The sensors are numbered 1-6 (top shelf, left to right), 7-12 (2^nd^ shelf), 13-18 (3^rd^ shelf), 19-24 (bottom shelf). The recording interface matches this layout as though you are standing in front of the rack looking at it.
b. Once you have placed the cage on the rack, move the large water sipper out of the way. Now, place the sipper holder on the cage lid (**Figure 10**).
c. Ensure that you have placed the correct sipper holder by checking that the number written on tape wrapped around the attached wires matches the sensor number for that animal

**Figure 10.**
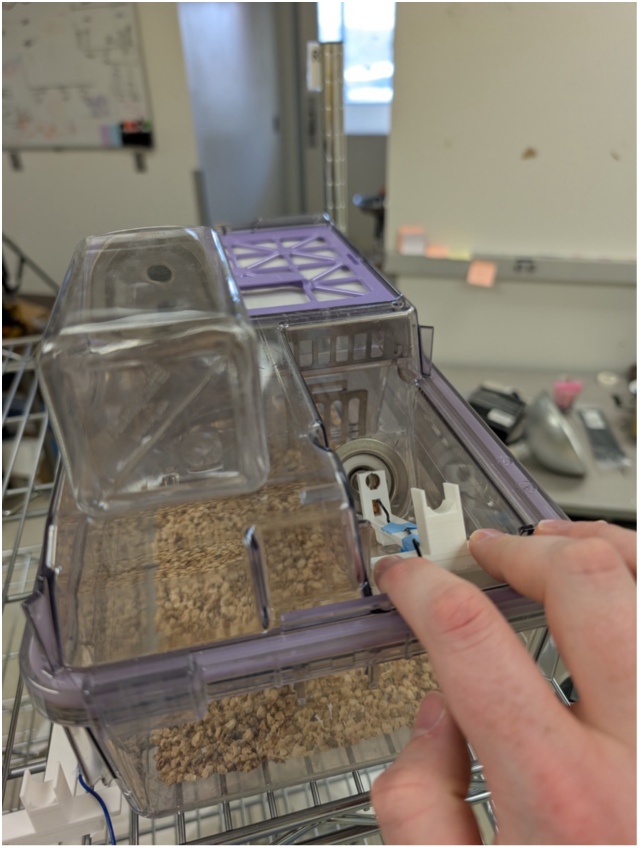
Place the sipper holder on the cage, moving the large water sipper out of the way.

O8. Once the cages are situated, press the “Start Recording” button. The sensor grid will change to look like **Figure 11** (note the colored start and test buttons).

**Figure 11.**
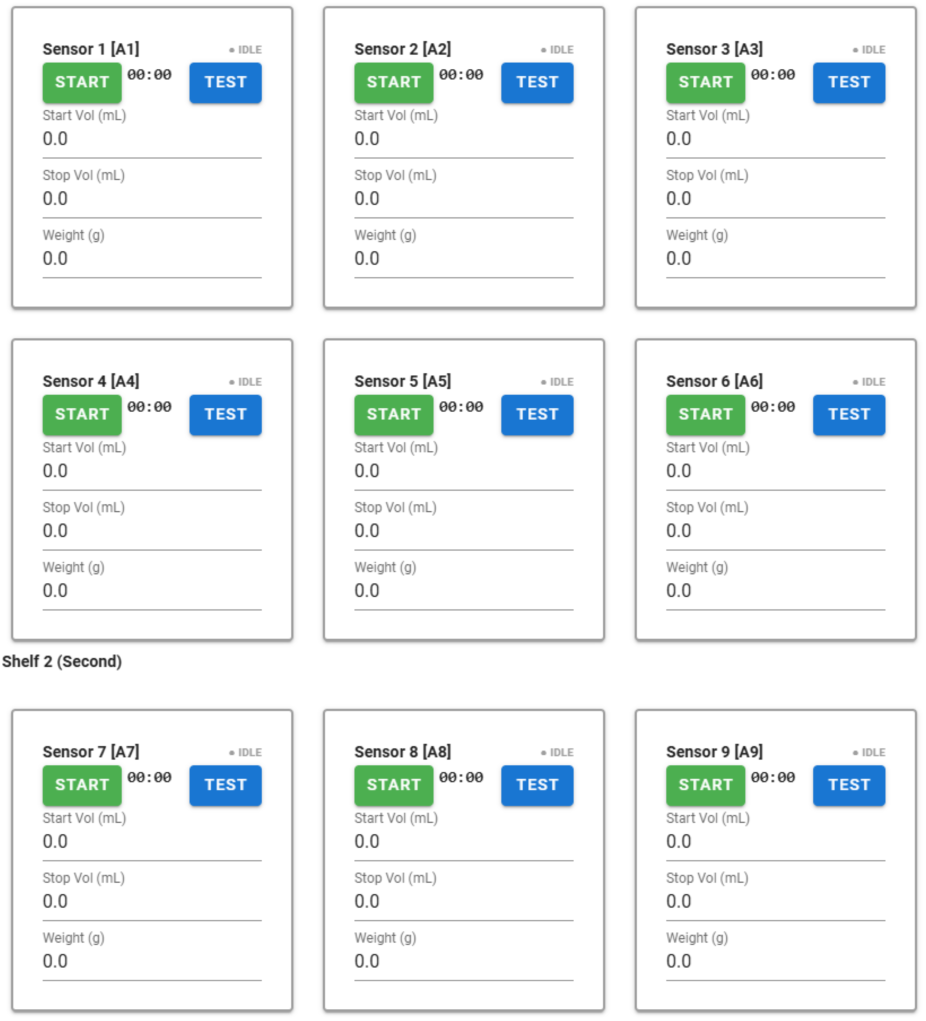
Sensor grid while recording, note the colored buttons (the buttons are greyed out when not recording).

O9. Starting with sensor 1, place the sippers in the holders one at a time while testing that the signal is good and recording volumes

a. Insert the sipper tip through the ring on the sipper holder and the metal opening on the cage lid.
b. Test the signal

i. Three firm presses with an ungloved finger on the metal ring through which we push the sipper
ii. Wait a second (for the data to be saved), then press the “Test” button next to the sensor you’re testing in the UI (**Figure 12**).
iii. You should be able to clearly distinguish all three finger presses
iv. If the signal isn’t good, there are a couple of things to try:

1. Use a small amount of alcohol to wipe the copper tape of the sipper holder and the sipper tip where it meets the holder (dirtiness can degrade the signal)
2. If you cannot get the signal to look good through fiddling, make a note of which sensor/animal in the comments box at the bottom of the CLiQR GUI and move on
c. Press the “Start” button corresponding to the sensor you just tested
d. Record the sipper volume in the “Start Vol” input box. See **Figure 13**. The volume should be read by the same person each time, both when starting and stopping a run, to minimize inter-person variability of readings. The sippers should typically be filled to around 10 mL, but the change in volume between starting and stopping volumes is much more important than the absolute starting point (a start reading of 2 mL and stop at 1.9 mL will give the same results as a start at 10 mL and stop at 9.9 mL in our analysis). Ensure that volumes are read to the nearest 0.05 mL (the best resolution we can read with graduation markings at 0.1 mL intervals).

**Figure 12.**
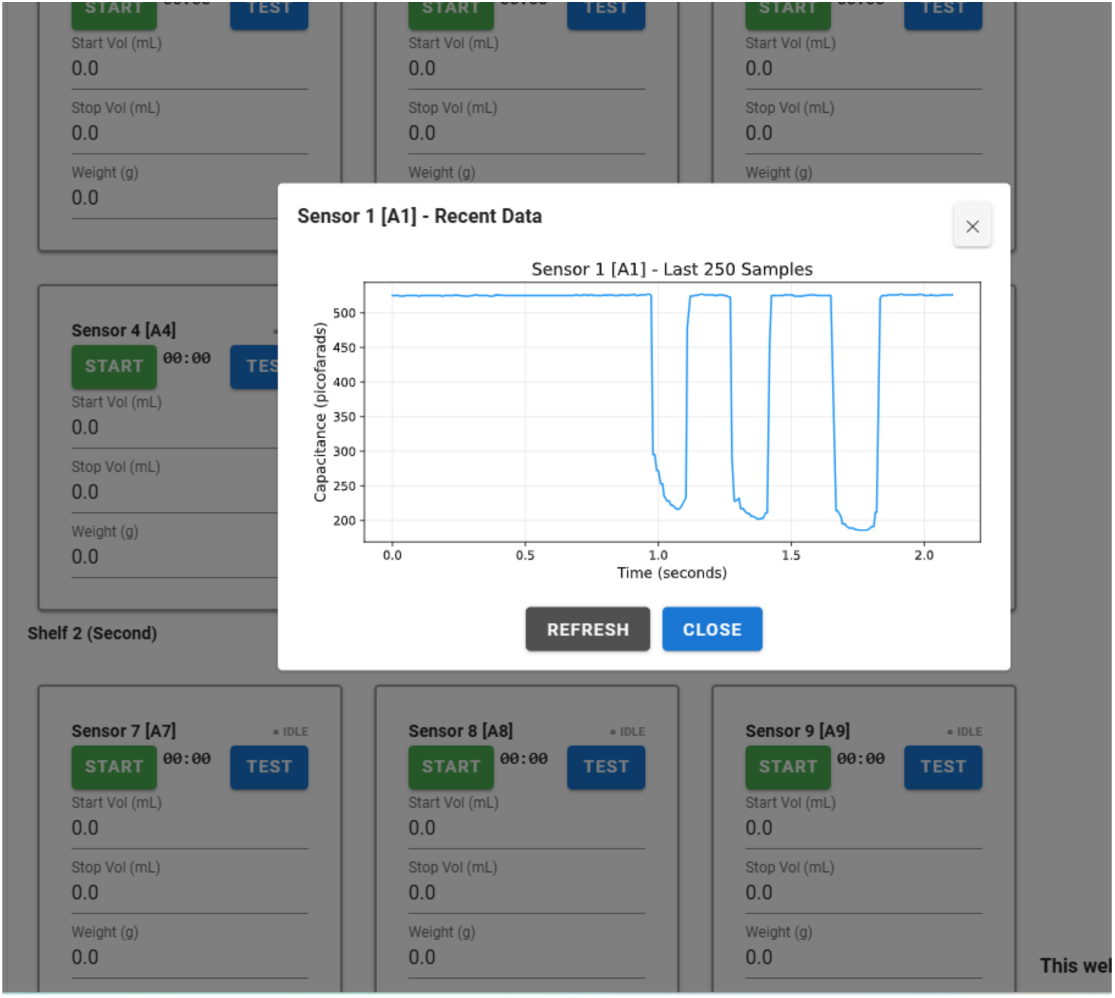
This is an example of a good test that shows a strong signal.

**Figure 13.**
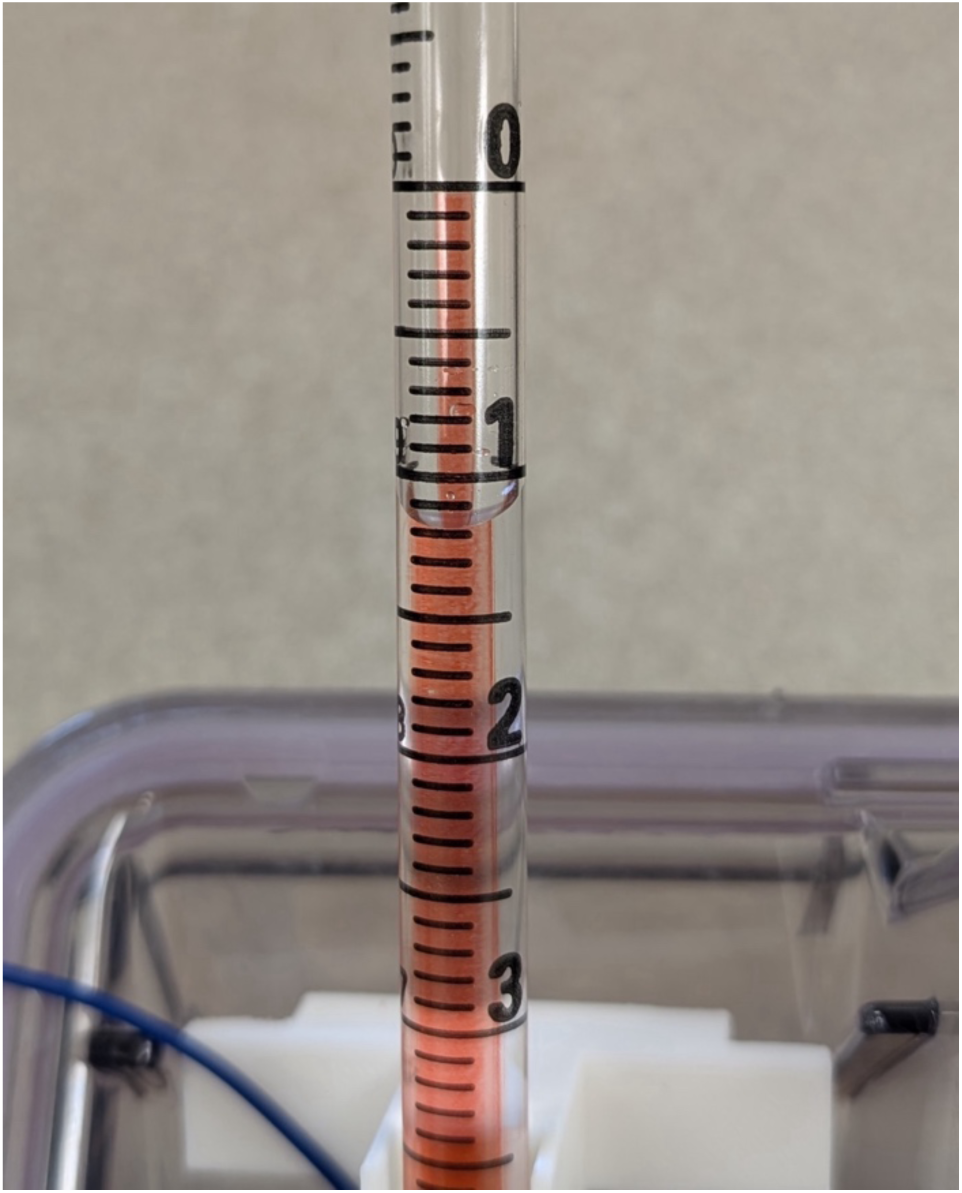
This sipper would be read as 8.8 mL. It’s difficult to see here, but there are alternative labels: 0 = 10, 1 = 9, 2 = 8, etc. So we read this as 1.2 mL below 10 = 8.8 mL.

O10. Once you have completed the above steps, set a timer for 2 hours

O11. Place the “Experiment in Progress” sign on the rack O12. Wait outside vivarium until the 2 hours are complete O13. Take “Experiment in Progress” sign down

O14. Starting with sensor 1 again:

a. Press the “Stop” button corresponding to the correct sensor
b. Read and record the volume of liquid remaining in the sipper in the “Stop Vol” input box
c. Carefully remove the sipper from the sipper holder and set aside
d. Remove the sipper holder from the cage lid and place it on the rack beside the cage
e. Replace the large water bottle
f. Transfer the cage back to the other rack

O15. Once you’ve stopped the sensors, press the large “Stop Recording” button to finalize the data files

O16. The data has been saved in the directory you selected for the “Output Directory”, which defaults to CLiQR\Lickometry Data\

a. raw_data_<date of recording>.h5

O17. Close the browser window and the PowerShell window (should be running behind the browser), then put the computer to sleep

O18. Empty the sippers, discarding remaining fluid

The “Comments” box found at the very bottom of the GUI window is a good place to make any notes about issues with the experiment. Any text entered here will be saved in the raw data HDF5 file and reported during the data analysis process.

## 7. Validation and characterization

### Key Capabilities and Limitations of CLiQR

- Strong positive correlation between detected licks and volume consumed, with R^2^ of 0.8494
- Solid precision (>98%) and recall (>94%)
- User-friendly GUI for recording sessions and data analysis
- Adaptability to various contexts in preclinical research beyond alcohol
- Currently only supports short-access experimental designs due to sipper volume limitations

### Validation

All animal procedures were approved by the University of Cincinnati Institutional Animal Care and Use Committee. We gathered lickometry data from 8 C57BL/6J mice completing six days of 2-hour drinking-in-the-dark (DID) sessions[26–28] (3 animal-sessions were not completed, given a total of 45 session recordings).

Mice were matched by sex, with 4 female and 4 male, and the DID sessions started at approximately 18 weeks of age. Sex was observed to have the expected effects (males weighed more and thus drank more: 0.461 ± 0.059 mL for males, 0.244 ± 0.104 mL for females, p = 0.029), but no effect of sex was observed in bout characterization metrics such as interlick intervals (123.0 ± 4.6 ms for males, 124.2 ± 8.5 ms for females, p = 1.0) and bout durations (5.96 ± 2.45 ms for males, 3.66 ± 1.33 ms for females, p = 0.114). **Figure 14** shows an overview of the recorded data, including example readings from the MPR121 boards and a simple linear regression of number of licks versus volume consumed.

**Figure 14.**
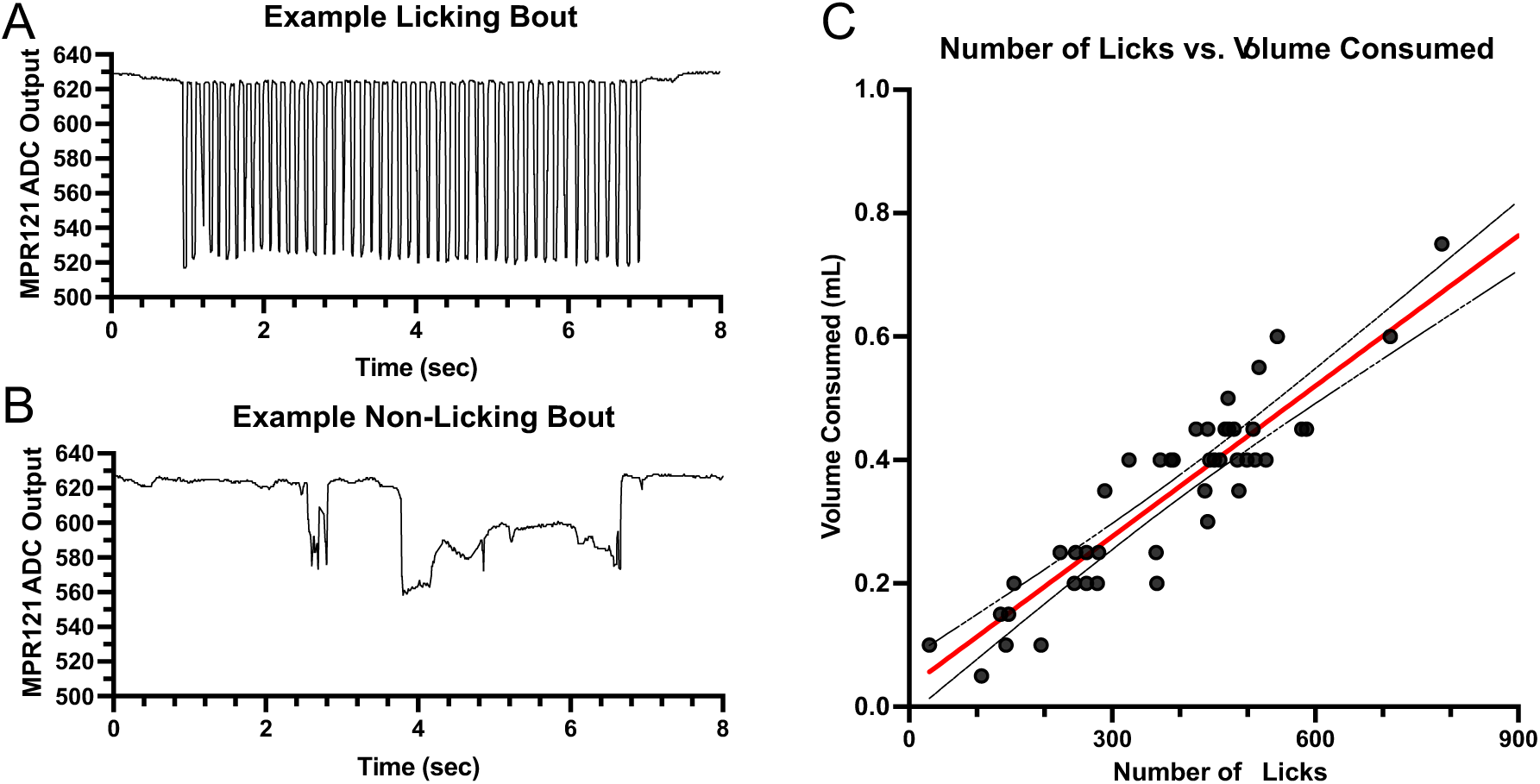
A) An example of the capacitive data recorded during a bout of licking. There is some variability in the capacitance values recorded during licks, due to variability in the contact of the tongue with the sipper. B) An example of the capacitive data recorded from a mouse climbing on top of the sipper. Rather than sharp peaks in the data, we see the capacitance values suppressed from baseline on the order of seconds, rather than milliseconds. C) Simple linear regression between number of licks and total volume consumed shows a strong positive relationship, R2 = 0.8494, p < 0.0001.

During periods that likely represent licking (**Figure 14A**), the signal exhibits rapid, millisecond-scale downward deflections from baseline. While the amplitude of individual deflections varies (probably due to subtle differences in contact geometry and moisture), the temporal structure remains sharply defined. In contrast, non-licking interactions such as climbing or resting on the sipper (**Figure 14B**) produce slower, broader shifts in capacitance on the order of seconds. These low-frequency changes are visually and algorithmically separable from candidate lick events.

To validate that detected events reflect fluid consumption, total lick counts were compared with volume consumed for each drinking session and animal. The pooled data showed a clear positive relationship between licks and intake (**Figure 14C**) when a simple linear regression is applied. The regression gave a result of R^2^ = 0.8494 (95% CI [0.7728, 0.9260]), with slope estimated to be 0.0008129 mL per lick (95% CI [0.0007076, 0.0009182]). This result confirms that the number of licking events detected from the capacitive signal provides a proxy for total consumption across many individual trials.

As shown in **Figure 15**, we report various descriptive statistics that characterize the detected licking behavior. For these analyses, we have grouped licking events into “bouts” of licks. The definition of bout we use follows that of other researchers utilizing C57 mice[1,2]: >1000 ms interlick interval (ILI) ends a bout of licking, and bouts must contain more than a single lick. Within-bout ILIs showed a strongly right-skewed distribution, with a median of 120.8 ms (corresponding to a licking frequency of 8.3 Hz) but a mean of 133.6 ms (7.5 Hz) (**Figure 15A**). A similar pattern emerged for licks per bout (median = 26, mean = 33.7) (**Figure 15B**) and bout duration (median = 3.25 s, mean = 4.37 s) (**Figure 15C**). In each case, the substantial discrepancy between median and mean values reflects the influence of a relatively small number of bouts with unusually long durations and high lick counts, which inflate the means while leaving the medians (which are more representative of typical bout structure) largely unaffected. Examination of licking dynamics across the session revealed a consistent pattern: animals exhibited modest front-loading during the initial portion of the session, followed by a substantial decrease in consumption rate, before settling into a relatively steady rate of consumption after approximately 40 minutes (**Figure 15D**). The cumulative bout count followed a similar trajectory (**Figure 15E**), indicating that changes in licking output across the session were driven by changes in bout initiation rather than by alterations in within-bout structure.

**Figure 15.**
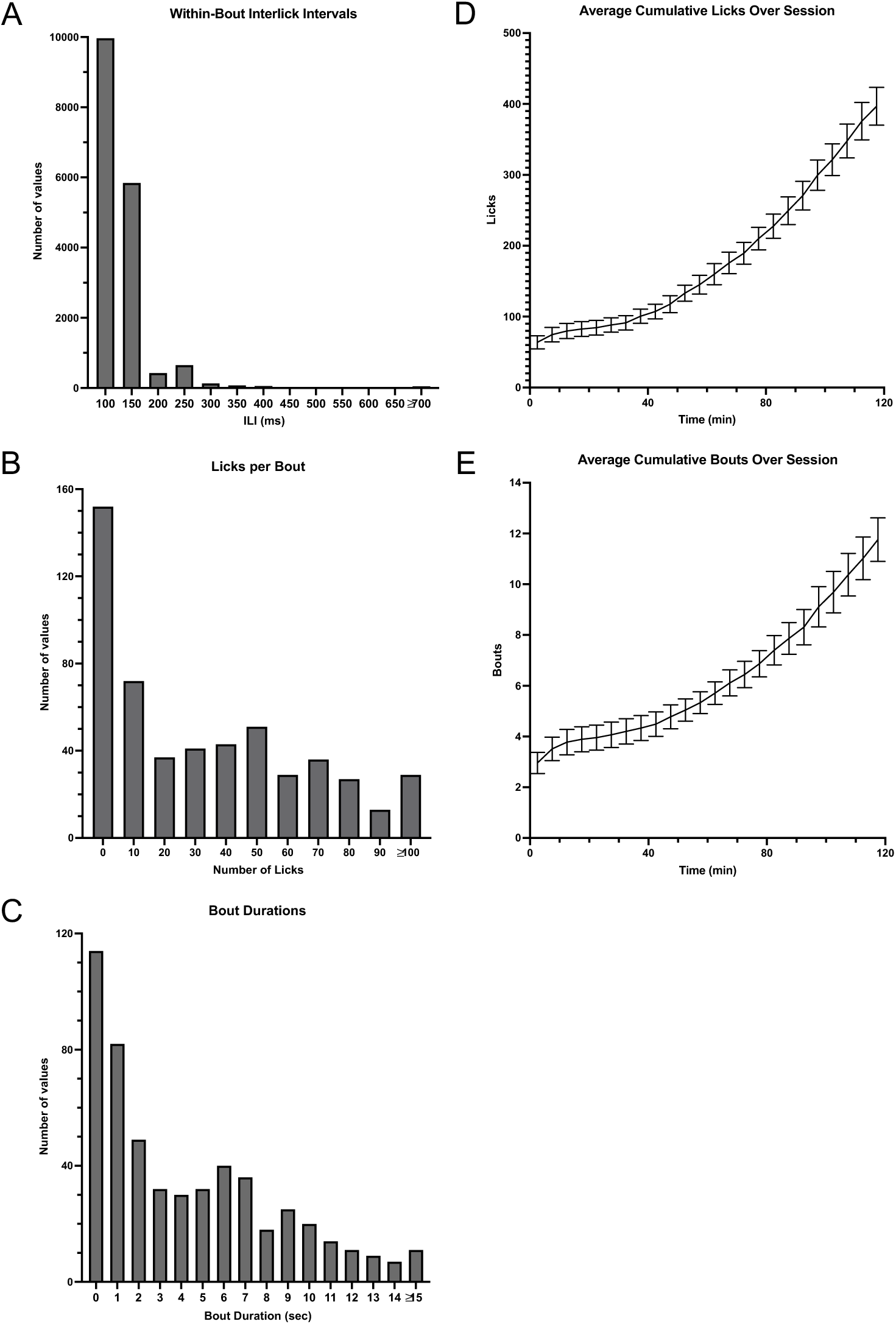
We used the following definition of bout: bouts must include 2 or more licks, with licks separated by more than 1000 ms ending a bout[1,2]. A) Histogram of within-bout interlick intervals (ILIs). The median ILI was 120. ms (8.3 Hz licking) and the mean ILI was 133.6 ms (7.5 Hz licking). Maximum within-bout ILI was capped at 1000 ms, and the small number of ILIs greater than or equal to 700ms have been grouped on the graph. B) Histogram of licks recorded per bout. A small number of bouts had greater than or equal to 100 licks and have been grouped together to make the graph more clear. The median licks per bout was 26 and the mean was 33.7. C) Histogram of bout durations. A small number of bouts had durations greater than or equal to 15 seconds, and these have been grouped together on the histogram. The median bout duration was 3.25 s and the mean was 4.37 s. D) Average cumulative licks over session. This graph shows the mean ± standard error across all animals and sessions (with licks binned in 5-minute chunks). We see a slight frontloading followed by a decrease in the rate of licking, then after ∼40 minutes the mice continue licking at a relatively consistent rate. E) Average cumulative bouts over session. This graph mirrors the graph in part D except that it shows the number of bouts instead of licks. The trend is similar to that in part D.

Quantification of false positive and false negative rates required concurrent video validation. We implemented a concurrent video recording feature in the CLiQR GUI that interfaces with a Raspberry Pi 5 running a Pi Camera Module 3 focused on the region of the cage near the sipper (no IR filter, used with IR floodlight for recordings in the dark). The system records at 120 frames per second, storing an exact time stamp for each frame to enable the sync. This feature automatically syncs the capacitance and video data and can be used to quickly generate clips for review/validation. We recorded six 2-hour DID sessions and we utilized DeepLabCut[29] to identify segments of video where the mouse was near the sipper for annotation of licking bouts. Across the 12 hours of video, CLiQR detected 3096 licks using the capacitance recording, with 3065 of these occurring inside of a licking bout annotated in the videos. By assuming that any detection outside of an annotated bout is a false positive, we obtain an estimated false discovery rate of 1.001% (95% CI [0.706, 1.418]). We have also estimated the false negative rate by analysis of the videos. In this procedure, we selected all drinking bouts where the mouse drank such that all tongue extensions were easily visible. By hand-scoring the total number of visible tongue extensions in the videos and comparing to the number of detected licks, we obtain an estimate of the number of false negatives. It should be noted here that we did not detect a single false positive during these bouts, leading to our estimate above. We stress that this is a rough estimate that includes a fair bit of uncertainty in scoring tongue extensions (mice sometimes partially extend the tongue without contacting the sipper). Based on whether we scored ambiguous partial extensions as full licks, we present two estimates of the false negative rate: 1.126% (527 total detections, 533 tongue extensions, 95% CI [0.23, 2.02]) or 4.356% (527 total detections, 551 tongue extensions, 95% CI [2.65, 6.06]). We believe that the true false negative rate falls somewhere between 0.23% and 6.06%. This equates to precision of 98.99% (95% CI [98.58, 99.29]) and recall of 98.87% (95% CI [97.98, 99.77]) or 95.64% (95% CI [93.94, 97.35]), with recall likely falling between 93.94% and 99.77%.

The synced video clips of all times where DeepLabCut detected the mouse in frame are available on Mendeley Data in the “All Clips with Mouse Visible” directory (https://data.mendeley.com/datasets/wpskw7k3mt/1). We have also extracted a few representative examples of licking bouts and climbing, and these are included in the “Curated Clips” directory of the Mendeley Data repository. All videos were uploaded with the metadata set to 30 fps, resulting in 25% default playback speed (given the 120 fps of the recordings).

### Characterization

The accurate quantification of rodent licking behavior is a necessity in various neuroscience and basic physiology studies. The choice of which lickometry system to use is heavily dependent on the experiments to be run and available funding. For our lab, the necessity of recording from dozens of mice concurrently while saving the full time series of data and keeping costs low motivated our creation of the CLiQR system.

Using a central small-form-factor desktop computer mounted to the housing rack, we record from up to 24 animals concurrently (including potential for 2-bottle choice experiments). By mounting the desktop directly to the rack with a touchscreen monitor and keyboard, we can present the user with an easy-to-use GUI, while also providing ample storage capacity for many data recordings. We believe that the addition of a full-size touchscreen GUI is a considerable improvement over many existing systems. This helps the laboratory personnel performing the experiments easily and helps decrease human error in recording weights/volumes in a darkened room. The system is limited by the 10 mL capacity of our custom sippers to short access paradigms, but future versions of the system will expand the system to function in longer access paradigms.

Our 2-hour DID sessions produce small total consumption volumes (∼1 mL), so the 0.05 mL measurement resolution in our manual volume readings represents a meaningful fraction of total intake. As such, the reading of volumes must be done carefully and consistently. Also, lab personnel must check carefully for leaking sippers as leaks can significantly bias results. We anticipate that R² values between number of licks and volume consumed will improve with more total data and when the system is used with longer-access paradigms and larger sipper reservoirs.

Future upgrades to the CLiQR system will include wireless capacitive sensors, automated volume recording with machine vision, and additional 3D print files for other types of cages and bottles. By saving the full data along with processed data, we allow future improvements in the algorithm used for lick classification to be applied retroactively. Increasing the size of the fluid reservoirs will additionally allow for adaptation of the system to research with rats. Finally, while we believe that the videos, and analysis thereof, we have demonstrated the proper functioning of the system, we plan to capture additional concurrent video and capacitive data to allow for yet more robust validation of lick detection in short-access paradigms.

In this paper, we have presented the Capacitive Lick Quantification in Rodents (CLiQR) system and the motivations for its creation. The data gathered with the system to date shows a strong correlation between volume of alcohol consumed and number of licks detected, although we believe there is room for improvement. The system was designed with future-proofing in mind, and we believe that it can be easily adapted for a variety of experimental contexts.

## Ethics statements

All animal experiments described in this manuscript were carried out under the oversight of the University of Cincinnati Institutional Animal Care and Use Committee, in accordance with the NIH Guide for the Care and Use of Laboratory Animals. The sex of animals was both male and female, with matched N, and did not significantly impact results. We have followed the ARRIVE reporting guidelines in writing this manuscript. Due to the nature of the experiments being for system validation, rather than testing a scientific hypothesis, we utilized already available animals, which limited the maximum N and sex makeup. This decision was made to reduce the total number of animals sacrificed in our lab. We also did not perform any control group tests or blinding/randomization.

## CRediT author statement

**Christopher Parker:** Conceptualization, Data curation, Formal analysis, Investigation, Methodology, Software, Validation, Visualization, Writing – Original draft, Writing – review and editing

**Alexis Lam:** Data curation, Investigation, Methodology

**Ashley Walters:** Investigation

**Harrison Carvour:** Investigation

**Jack Douglass:** Investigation

**Brannen Dyer:** Investigation

**Anthony Glorius:** Investigation

**Boston Main:** Investigation

**Christine Moore:** Investigation

**Margaret Niemeier:** Investigation

**Avni Patel:** Investigation

**Kaladin White:** Investigation

**Nicholas Timme:** Conceptualization, Funding Acquisition, Methodology, Project administration, Resources, Supervision, Writing – review and editing

## Acknowledgments

This work was supported by the National Institute on Alcohol Abuse and Alcoholism of the National Institutes of Health: grant # R00AA028265 to NT

## Declaration of generative AI and AI-assisted technologies in the manuscript preparation process

Anthropic’s Claude Opus 4.8 was used to assist in drafting the Abstract section of this manuscript.

## Notes

https://github.com/TimmeLab/CLiQR

### Competing Interest Statement

The authors have declared no competing interest.

### Summary of Updates

Updated to the HardwareX format, and additional validation experiments and system improvements were carried out.

